# A mechanistic path to maximize biomass productivity while maintaining diversity

**DOI:** 10.1101/800706

**Authors:** Oscar Godoy, Lorena Gómez-Aparicio, Luis Matías, Ignacio M. Pérez-Ramos, Eric Allan

## Abstract

With ongoing biodiversity loss, it is important to understand how the mechanisms that promote coexistence relate to those that increase functioning in diverse communities. Both coexistence and biodiversity functioning research have unified their mechanisms into two classes. However, despite seeming similarities, theory suggests that coexistence and biodiversity mechanisms do not necessarily map onto each other, yet direct empirical evidence for this prediction is lacking. We coupled field-parameterized models of competition between 10 plants with a biodiversity-functioning experiment measuring biomass production, litter decomposition, and soil nutrient content under contrasting environmental conditions. We related biodiversity mechanisms (complementarity and selection effects), to coexistence mechanisms (niche and fitness differences). As predicted by theory, complementarity effects were positively correlated with niche differences and differences in selection effects were correlated with fitness differences. However, we also found that niche differences contributed to selection effects and fitness differences to complementarity effects. Despite this complexity more stably coexisting communities (i.e. those in which niche differences offset fitness differences) produced more biomass, particularly under drought. This relationship was weaker for litter decomposition rates and soil nutrient acquisition, showing that the mechanisms promoting plant coexistence may differ from those promoting high levels of functions that are less directly related to plant performance. We provide the first empirical evidence that the mechanisms promoting stable coexistence correlate with those driving high biomass production. These findings establish a link between stable coexistence and functioning, which could allow better predictions of how diversity loss induced by global change translates to changes in ecosystem function.

## Introduction

A large number of experimental and observational studies have shown that more diverse communities tend to have higher levels of multiple ecosystem functions (1–5). At the same time, global change is reducing opportunities for coexistence (6–7), and therefore diversity (8), ultimately reducing a range of ecosystem functions and services (9). A better understanding of the mechanistic link between the processes maintaining biodiversity in communities and those driving functioning would allow us to better predict effects of global change on ecosystem functioning and to optimize restoration efforts. However, links between the conditions necessary for stable coexistence and the processes driving high functioning in diverse communities have remained elusive.

Coexistence and biodiversity-functioning theory have both developed frameworks unifying their principle mechanisms into two groups. Two main classes of mechanisms have been identified as underlying effects of biodiversity on function (10): complementarity effects occur when species, on average, yield more in mixture than in monoculture, and selection effects occur when there is a covariance (positive or negative) between monoculture yield and dominance in mixture. Similarly, modern coexistence theory (11) has successfully grouped coexistence mechanisms into: stabilizing processes which enhance niche differences, through increasing intraspecific competition relative to interspecific competition, and equalizing processes, where fitness differences that lead to competitive dominance are minimized. A first step towards combining biodiversity-functioning and coexistence theory would therefore be to show how these classes of mechanisms relate to each other. This would also clearly demonstrate how the causes of complementarity link to its effects, which has been a major source of confusion (12). There is a general assumption that niche differences, for instance in how species exploit resources, drive positive complementarity effects (10, 13–15) and therefore that strong complementarity between species should allow them to coexist. It is also tempting to assume that differences in fitness between species, i.e. in competitive dominance (11), produce selection effects. However, recent theoretical studies have shown that these mechanisms may not relate directly and that in fact positive complementarity effects can occur even without stabilizing niche differences (16–17). Thus, it is not clear to what extent biodiversity and coexistence mechanisms relate in practice. Surprisingly, while a huge number of studies have quantified selection and complementarity, no empirical studies have tried to link them explicitly to measured coexistence mechanisms.

Recent theory shows that the relationship between selection and complementarity effects and niche and fitness differences can be more complicated than initially assumed (15, 18). The main reason is that selection and complementarity effects are determined by species’ relative abundances and density dependent effects, and these emerge from the combination of niche and fitness differences between the species (19–21). Theoretical advances therefore suggest that we need to know what combination of niche and fitness differences maximize functioning. With larger niche differences promoting species evenness and smaller fitness differences reducing competitor dominance, our main hypothesis is that the communities with the highest functioning will be those in which the stabilizing effect of niche differences is sufficient to offset the fitness differences between species and therefore to allow them to stably coexist. Selection and complementarity effects have almost always been assessed for biomass production, but other critical ecosystem functions may show different responses to biodiversity (1). We therefore also aim to test whether the conditions promoting stable coexistence also promote high levels of functions other than biomass production such as litter decomposition or soil nutrient cycling.

To rigorously evaluate the relationships between biodiversity-functioning and coexistence mechanisms, we performed a combined competition and biodiversity functioning experiment with 10 annual plant species (Table 1) in a Mediterranean grassland. We field-parameterized population models to quantify stabilizing niche differences and average fitness differences. Then, we related these pairwise species differences to observed complementarity and selection effects for biomass, litter decomposition and soil nutrient variation. Prior experimental and observational work has shown that environmental variation modulates the net effect of diversity on productivity (22, 23), which is consistent with theoretical expectations (24). To further evaluate the role of environmental variation, we directly manipulated the timing of species germination to create two contrasting scenarios of water availability (control climate and drought treatments; see the section “Methods” for more details). This manipulation modified the niche and fitness differences between species pairs (7) and allowed us to test for the strength of the correlation between niche and fitness differences and complementarity and selection effects in different environments.

**Table 1.**
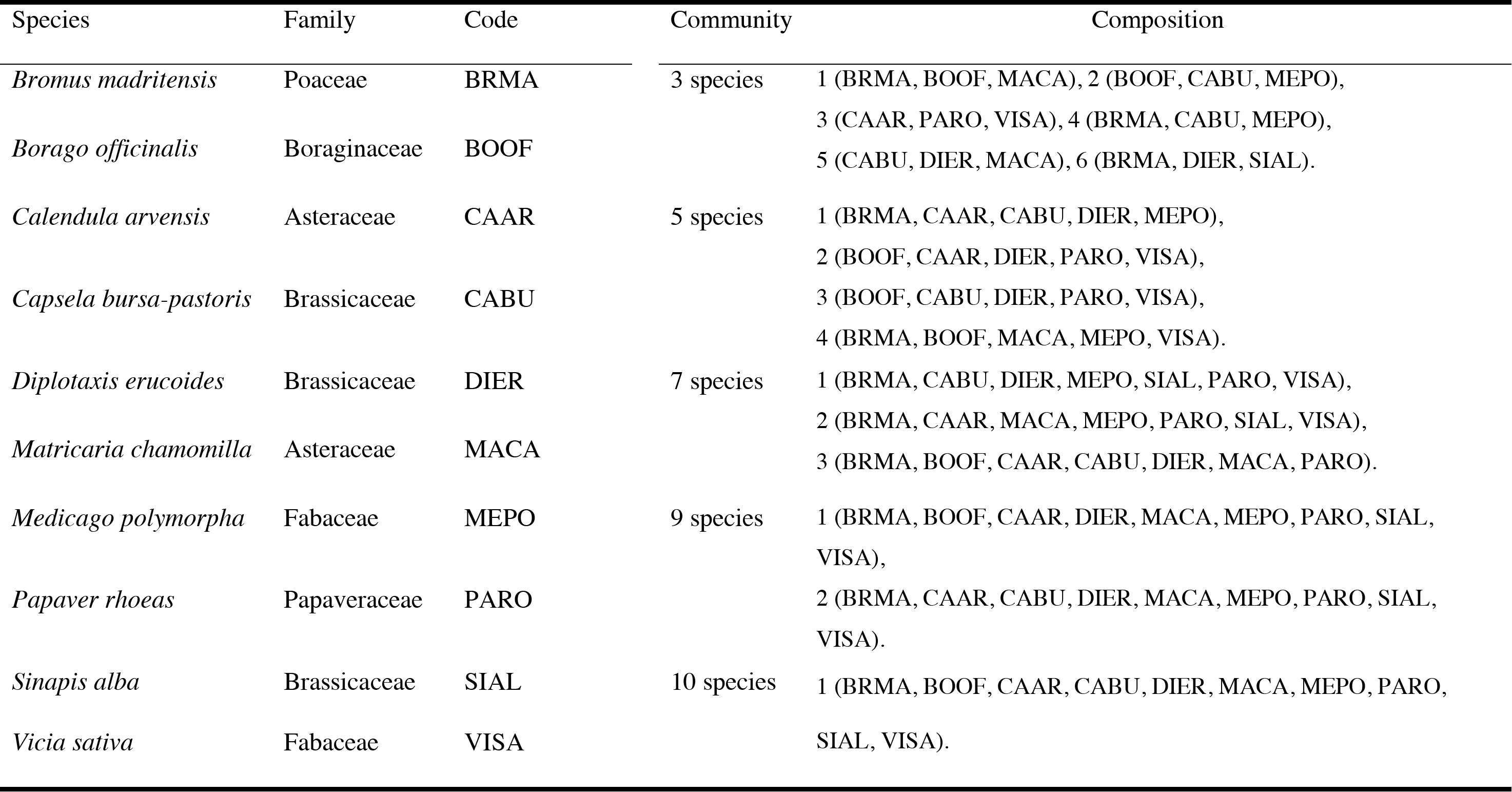
Species and assembled communities with the diversity levels used in the experiment.

## Results

We first analyzed the overall relationship between diversity and function and found that more diverse communities produced more biomass, their litter decomposed faster and showed lower soil N than less diverse communities, although the magnitude of these relationships depended on the climatic conditions (Fig. 1). These positive diversity effects mostly resulted from an increase in complementarity effects with increasing community diversity. In contrast, selection effects became more negative with increasing diversity (*SI Appendix*, Fig. S1).

**Fig. 1.**
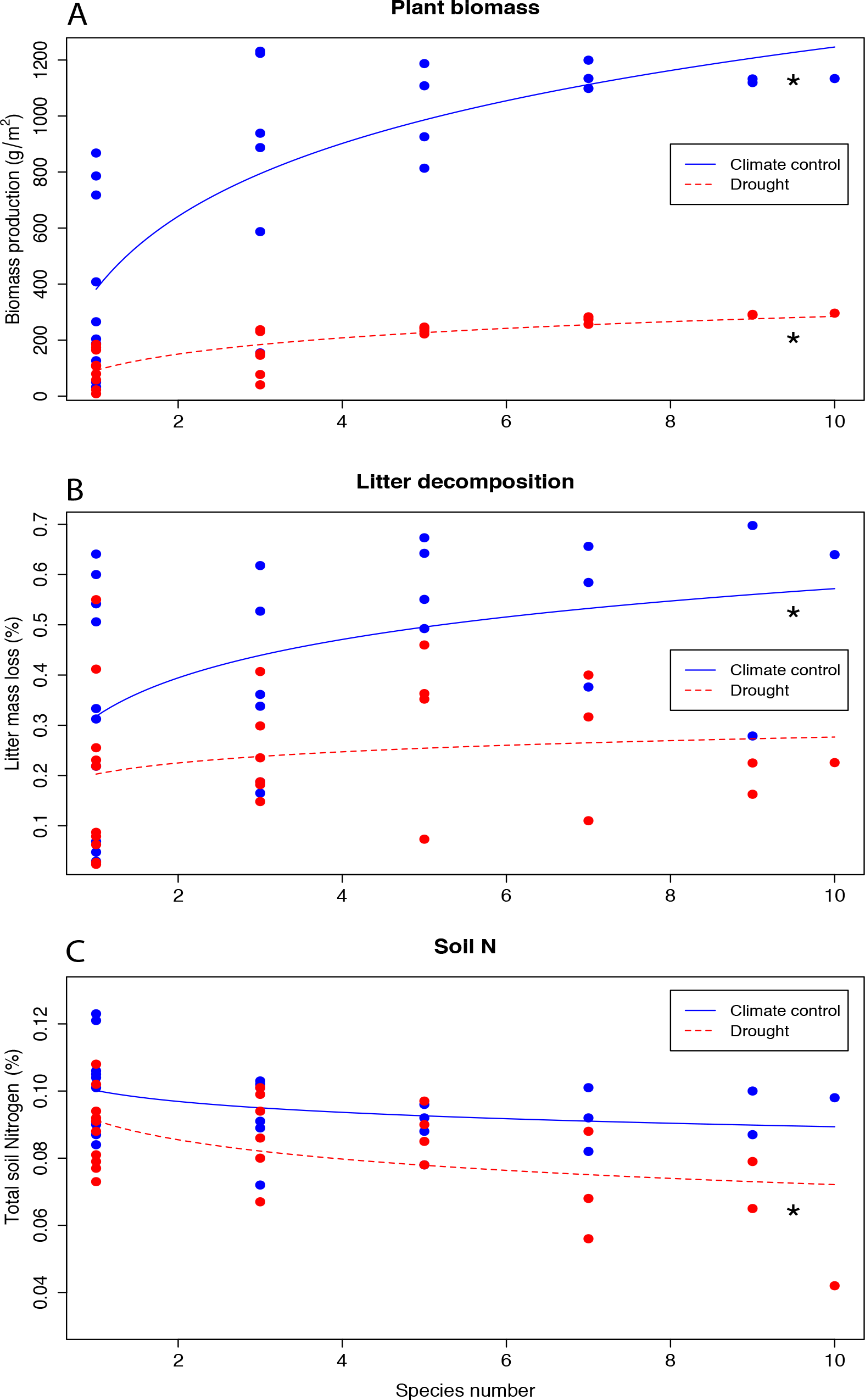
Observed effects of species diversity on biomass production, litter decomposition and soil nutrient availability. Total Soil N is shown here as an example of soil nutrient content but very similar relationships were observed for the other soil elements measured (C, P, Ca, Mg, and K). Blue lines and points represent the communities under control climate conditions and red dashed lines and points show the same communities under drought conditions. Non-linear instead regressions fitted the data better. Significant regressions at P < 0.05 are represented with an asterisk.

We next related biodiversity-functioning to coexistence mechanisms. Niche and fitness differences are defined for pairs of species (15) in the annual plant model (see eq. 2 and 3, material and methods), but complementarity and selection effects are commonly measured at the community level (10). To compare the effects, we therefore selected techniques (diversity interaction models (25)) that allowed us to calculate complementarity between pairs of species and selection effects for individual species, which we converted to pairwise differences in selection effects. When averaged at the community level, these selection and complementarity effects were similar to those obtained from the additive partition of Loreau & Hector (10) (*SI Appendix*, Fig. S2). For all functions evaluated, pairwise complementarity effects were higher when stabilizing niche differences were large and when average fitness differences were small. In contrast, pairwise differences in selection effects were larger when niche differences were small and fitness differences were large (Fig. 2). Although the direction of effects was consistent across functions, their significance varied and complementarity effects were sometimes only weakly linked to niche differences (Fig. 2). Surprisingly, these relationships across functions were in general stronger under drought conditions, despite the fact that the drought treatment significantly reduced both niche and fitness differences (7) (*SI Appendix*, Fig. S3).

**Fig. 2.**
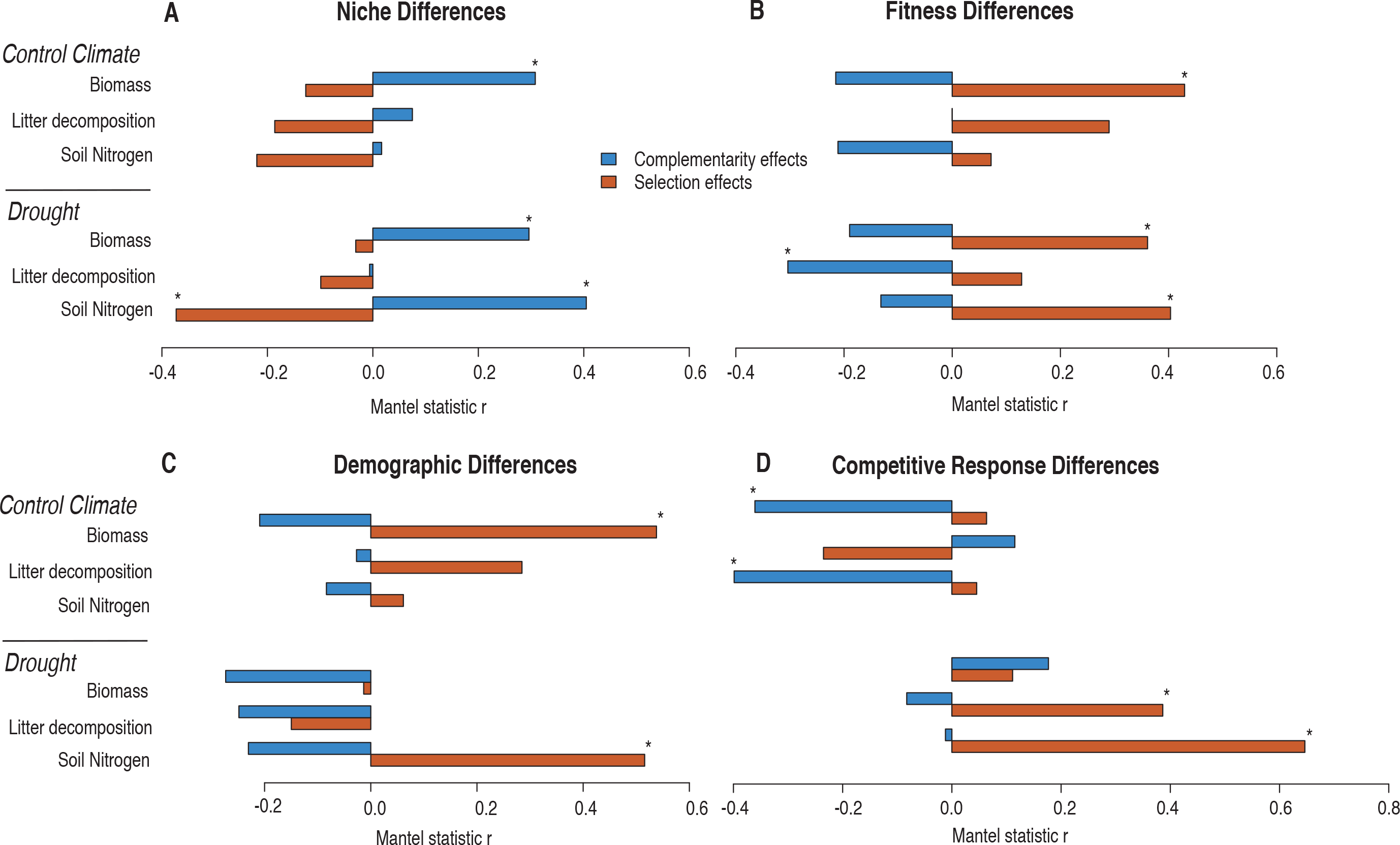
Correlations between (A) stabilizing niche and (B) average fitness differences and complementarity (blue) and selection effects (red). Correlations are shown for the three functions considered, under the two contrasting environmental conditions (control climate, drought). Correlations between complementarity and selection are also shown with the two components of fitness differences, the demographic ratio (C) and the competitive response ratio (D). Significant correlations, following Benjamini-Hochberg correction for multiple comparisons, are marked with an asterisk.

Fitness differences between species can result from differences in demography (i.e. differences in intrinsic growth rates) or in species responses to competition (see material and methods, eq. 3). In order to investigate the importance of these two components, we correlated each one with complementarity and selection and found that they did not contribute equally to observed complementarity and selection effects. Complementarity effects only correlated significantly with the competitive response ratio, not the demographic ratio, and only under the control climate (Fig. 2). This suggests that asymmetries in species’ sensitivity to competition, rather than differences in growth rate, reduced complementarity effects between them. In contrast, both demographic and competitive response differences were correlated with differences in species’ selection effects. Demographic differences correlated with differences in selection effects on biomass (control climate) and soil nitrogen content (drought) and competitive response differences correlated with differences in selection effects for soil nitrogen content and litter decomposition under drought (Fig. 2). Importantly, the relationships observed for soil N were the same for the other soil elements analyzed, namely total organic carbon, available P, and exchange cations (*SI Appendix*, Fig. S4).

Both stabilizing niche differences and average fitness differences influenced multiple ecosystem functions. We therefore evaluated their combined effect on functioning, i.e. whether species pairs predicted to more stably coexist have higher functioning. Supporting our main hypothesis, we found that the species pairs predicted to coexist more stably, i.e. those in which observed niche differences were larger than the minimum required to offset fitness differences, were in turn predicted to produce significantly more biomass under both climatic conditions. However, they were not predicted to have higher levels of the other functions, i.e. litter decomposition or soil resource use (Fig. 3).

**Fig. 3.**
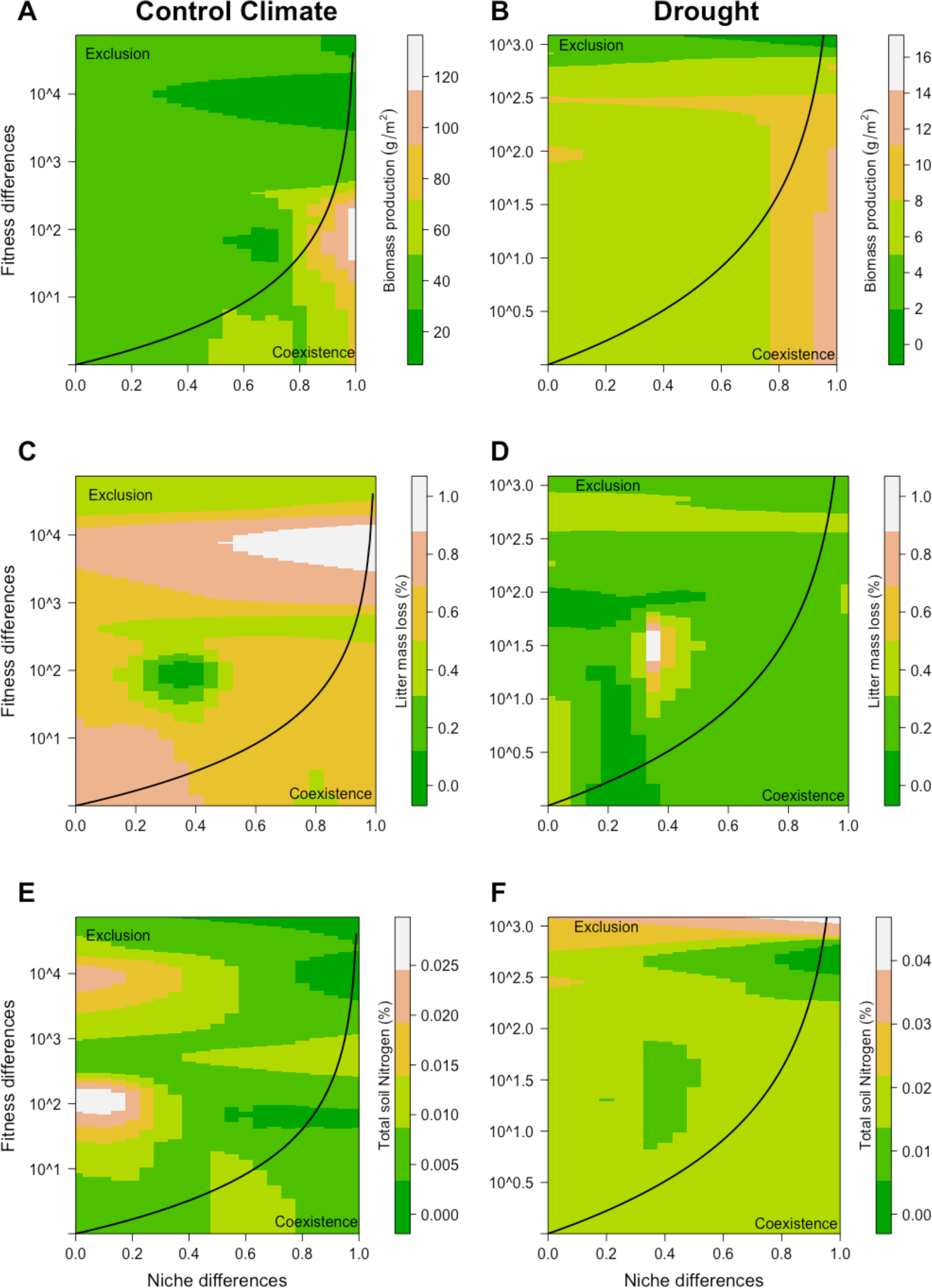
Relationship between the degree of stability of coexistence and the levels of multiple functions (biomass, upper two panels; litter decomposition, middle two panels; total soil N, lower two panels) under control climate conditions (left panels), and drought (right panels). The heat map represents the level of functioning estimated for a given species pairs, using the approach of (30). Greener colors represent low functioning while brownish to white colors represent higher functioning. The solid black line indicates whether the conditions for coexistence are met (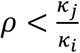, where species *j* is the fitness superior) and separates the coexistence from the competitive exclusion region. Mantel tests, following Benjamini-Hochberg correction for multiple comparisons, showed that species pair stability (i.e. their distance from the coexistence line) was significantly related to their predicted biomass production (Control conditions Mantel r = 0.38, P = 0.048; Drought Mantel r = 0.43, P = 0.018) but not to the predicted litter decomposition (Control conditions Mantel r = 0.14, P = 0.359; Drought Mantel r = − 0.26, P = 0.850) or soil nitrogen availability (Control conditions Mantel r = − 0.32, P =0.909; Drought Mantel r =−0.16, P =0.788). For a graphical representation of observed pairwise niche and fitness differences, see *SI Appendix*, Fig. S3.

To estimate the long-term stability of our experimental communities, we used the empirically estimated vital rates and interaction coefficients for all species pairs to build a competitive network. Diverse communities are locally stable only if all of the species can coexist despite perturbations to their population size at equilibrium. We did predict feasible and locally stable equilibrium for several species’ pairs and one triplet although not for any of the experimental assembled communities, under the two climatic conditions (*SI Appendix*, Fig. S5). This result suggests that observed levels of the ecosystem functions were transient rather than stable.

## Discussion

Understanding connections between the factors that promote species coexistence and high ecosystem functioning would allow a better mechanistic understanding of how biodiversity loss translates to reductions in ecosystem function. Both fields have developed frameworks to unify mechanisms but theoretical attempts to link the frameworks have shown that they cannot be easily mapped onto each other. However, by combining recent advances in coexistence theory with a series of competition and biodiversity functioning experiments, we could show that the two frameworks can be linked and that positive effects of biodiversity on functioning resulted from a combination of niche differences stabilizing coexistence, and average fitness differences driving competitive exclusion. Moreover, our results provide a clear link between the conditions for stable coexistence and high functioning by showing that biomass production is maximized when species coexist more stably, i.e. when niche differences exceed more strongly fitness differences.

According to the classic expectations, we provide empirical support for the assumption that niche differences drive complementarity, while competitive ability differences drive selection effects. However, our results also support recent theoretical suggestions (15–18) that both selection and complementarity effects include a combination of niche and fitness differences (Fig. 2). We generally find larger complementarity effects when species differ in their niches and larger differences in selection effects when species differ substantially in fitness. Depending on the function and environment, differences in selection effects could be driven either by differences in species intrinsic growth rates or differences in their response to competition. These results imply that communities should lead to functioning being driven by a smaller number of species when they contain species that vary more in growth rate, (e.g. differences among species in resource conservation versus acquisition) (26), or factors that enhance differences in competitive ability, such as high nutrient addition (27).

Complementarity effects were promoted by niche differences but were also reduced when species differed strongly in fitness, more specifically, if they differed in their response to competition. In addition, differences in selection effects between species were reduced when they differed strongly in their niches. The negative effects of fitness differences on complementarity, and the negative effects of niche differences on differences in selection effects likely arise because both niche and fitness differences are density dependent effects involving the same model parameters (see equations 2 and 3). This means that two interrelated processes should occur at the same time to increase function. First, fitting with the classic expectation, species should differ in their niches, but second and much less intuitive, species should have similar sensitivities to competition. Therefore, factors such as nutrient additions or loss of consumers that reduce niche differences or enhance differences in competitive ability are expected to reduce complementarity effects and to lead to a greater imbalance in species functional effects.

Given this more complex relationship between the determinants of coexistence and the mechanisms promoting positive effects of biodiversity on functioning, we also conducted an integrated analysis to determine if those communities where coexistence was most stable had highest functioning. In support of our main hypothesis, we found that more stable pairs (i.e. those in which their niche differences exceed more strongly their fitness differences) were predicted to produce more biomass (Fig. 3). Theory (15–18) shows that complementarity effects can arise even in the absence of stable coexistence; however, our empirical approach shows that this may be quite rare and that high biomass production is only likely in communities that can stably coexist. This is an important result because it provides a clear link between the conditions for stable coexistence and ecosystem functioning and therefore, a direct pathway to achieve the desired objective of maximizing functioning while maintaining biodiversity.

Although they were productive, more stably coexisting plant communities did not show faster litter decomposition rates or more soil nutrient uptake (Fig. 3). In addition, complementarity effects on litter were not explained by any of the coexistence mechanisms. These results suggest that the conditions leading to more stable coexistence of plant species maximize functions directly related to plant performance, such as biomass production, but have weaker effects on functions which are less directly related. A potential explanation of these mismatches is that these functions more strongly involve the effect of other trophic levels. For instance, litter decomposition is influenced not only by leaf litter traits, but also by the effect of soil organisms, including macro- and micro-invertebrates, nematodes, bacteria, and fungi (28). In addition, the combinations of plant traits that lead to high litter decomposition, or soil nutrient uptake, may be different to those determining stable coexistence. These results show that linking coexistence and functioning is likely to be more complex for functions other than biomass. To understand how diversity loss affects these functions we may need to consider the mechanisms promoting coexistence not only of plants but of multiple trophic levels (29).

Analyses based on single-stable equilibrium predicted that none of our experimental communities could coexist at the scale of the experiment (Table 1). This is not a surprising result since we could not include the spatial and temporal variations key to maintain diversity at larger scales (30), only coexistence mechanisms that operate in constant environments can contribute to the niche differences we measured (11). Nevertheless, we do find a significant link between stable coexistence and biomass production, which suggests that non-spatial/temporal coexistence mechanisms such as resource partitioning or natural enemies do promote high biomass production in this system. Evaluating how much function coexistence mechanisms operating in variable environments add to the system is definitely a promising direction for further research.

Environmental conditions affected the strength of relationships between coexistence and biodiversity mechanisms. In our study, delaying germination decreased rainfall by almost 40% and reduced the growing season by two months. This delay strongly reduced biomass production to about 10% of the level in control conditions (Fig. 1), consistent with the predominant role of water availability in controlling biomass yields in Mediterranean ecosystems (e.g. 31). This reduction in water availability might be expected to reduce available niches and competitive ability differences and we did find evidence that wetter environmental conditions allowed for greater niche and fitness differences between species pairs (*SI Appendix*, Fig. S3) (7). With greater niche overlap, it is reasonable to expect a weaker relationship between stabilizing niche differences and complementarity, however, we actually observed a stronger correlation between them (Fig. 2). This implies that not all of the processes driving niche differences contributed to complementarity effects under wet conditions (Fig. 2). Our approach is phenomenological, which means that we do not know the specific sources of variation in observed niche differences in our experiment. Overall, these results emphasize the context-dependency of biodiversity effects on functioning, and call for a framework to understand what type of environmental conditions are promoting the niche differences that contribute to complementarity effects.

Our study represents a step forward in evaluating the link between the drivers maintaining diversity and functioning compared to previous experimental work that considered particular components (e.g. interspecific facilitation, (32)) or aggregates (e.g. community evenness (33)) of niche and fitness differences. Still our approach to measure stability, either that of species pairs or of equilibrium communities, is fundamentally based on pairwise interactions between species. The next step is to move beyond this pairwise framework to one in which niche and fitness differences are estimated at the community-level, and in which additional factors determining coexistence at the multispecies level such as indirect or “higher order” interactions are incorporated (34). Although recent toolboxes have been proposed to adopt this new framework (35), we lack clear expectations about how the mechanisms determining the degree of stability in complex communities are linked to the net effect of biodiversity on functioning. However, incorporating these effects would be important to test if the communities providing high levels of functions like litter decomposition are in fact able to stably coexist. Multispecies interactions may be more important for explaining other functions or functioning in other contexts but our results suggest that, for biomass production at least, the degree of stability in coexistence predicted by the pairwise approach works well to explain functioning.

Here, we provide the first empirical evidence that niche differences do indeed drive complementarity effects. However, consistent with recent theoretical advances (15, 18), we found that both selection and complementarity are related to a combination of the stabilizing niche differences that promote species diversity and to the average fitness differences that promote competitive exclusion. Despite these complex relationships, when integrating the conditions for coexistence, we found that more stable coexistence promotes higher biomass production. This implies that any process that destabilize coexistence should therefore reduce ecosystem functioning. However, extending these findings to functions beyond biomass is more complex. Overall, our results suggest that we need to develop a framework to link the species differences allowing more stable coexistence to those promoting high levels of different functions. This would involve expanding our perspective to multitrophic communities and perhaps to indirect and higher order interactions. We also need to complement the phenomenological approaches used here with more detailed consideration of underlying mechanisms. Our results provide a first step in this direction by showing that the conditions promoting stable coexistence and high ecosystem functioning are the same.

## Material and Methods

### Study site and experimental setup

Our experiment was conducted at the *La Hampa* field station of the Spanish National Research Council (CSIC) in Seville, Spain (37°16’58.8”N, 6°03’58.4”W), 72 m above sea level. The climate is Mediterranean, with mild, wet winters and hot, dry summers. Soils are loamy with pH = 7.74, C/N = 8.70 and organic matter = 1.16% (0-10 cm depth). Precipitation totaled 532 mm during the experiment (September 2014-August 2015), similar to the 50-y average. We used ten common annual plants, which naturally co-occur at the study site, for the experiment. These species cover a wide phylogenetic and functional range and include members of six of the most abundant families in the Mediterranean grasslands of southern Spain (Table 1). Seeds were provided by a local supplier (Semillas silvestres S.L.) from populations located near to our study site. Our experiments were located within an 800 m^2^ area, which had been previously cleared of all vegetation and which was fenced to prevent mammal herbivory. Landscape fabric was placed between plots to prevent growth of weeds.

### Theoretical background for quantifying niche and fitness differences

Here we summarize the approach developed in (36) to quantify the stabilizing effect of niche differences and average fitness differences between any pair of species. Both these measures are derived from mathematical models that capture the dynamics of competing annual plant populations with a seed bank (21, 37). This approach has been used in the past to accurately predict competitive outcomes between annual plant species (36). Population growth is described as:

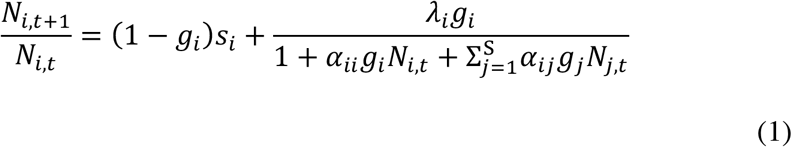

where 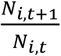 is the per capita population growth rate, and *N*_*i,t*_ is the number of individuals (seeds) of species *i* before germination in the fall of year *t*. Changes in per capita growth rates depend on the sum of two terms. The first describes the proportion of seeds that do not germinate (1 − *g*_*i*_) but survive in the seed soil bank (*s*_*i*_). The second term describes how much the per germinant fecundity, in the absence of competition (*λ*_*i*_), is reduced by the germinated density of conspecific (*g*_*i*_*N*_*i,t*_) and various heterospecific 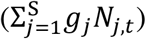 neighbors. These neighbor densities are modified by the interaction coefficients describing the per capita effect of species *j* on species *i* (*α*_*ij*_) and species *i* on itself (*α*_*ii*_).

Following earlier studies (11, 36), we define niche differences (1 − *ρ*) for this model of population dynamics between competing species as:

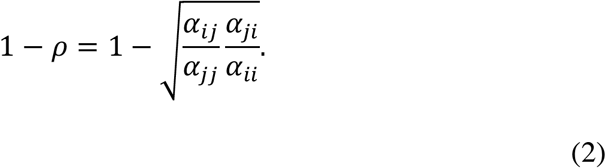

The stabilizing niche differences reflect the degree to which intraspecific competition exceeds interspecific competition. 1 − *ρ* is 1 when individuals only compete with conspecifics (i.e. there is no interspecific competition) and it is 0 when individuals compete equally with conspecifics and heterospecifics (i.e. intra and interspecific competition are equal). Niche differences between plant species can arise for instance from differences in light harvesting strategies (36, 38), or in soil resource use and shared mutualisms (39).

The average fitness differences between a pair of competitors is 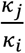 (36), and its expression is the following:

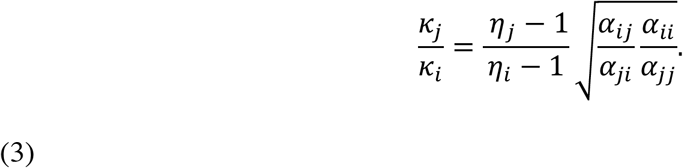

The species with the higher value of 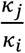 (either species *i* or species *j*) is the competitive dominant, and in the absence of niche differences excludes the inferior competitor. This expression shows that 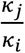 combines two fitness components, the “demographic ratio” 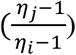 and the “competitive response ratio” 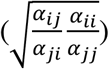. The demographic ratio is a density independent term and describes the degree to which species *j* has higher annual seed production, per seed lost from the seed bank due to death or germination, than species *i*:

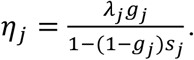

The competitive response ratio is a density dependent term, which describes the degree to which species *i* is more sensitive to both intra and interspecific competition than species *j*. Note that the same interaction coefficients defining niche differences are also involved in describing the competitive response ratio, although their arrangement is different.

With niche differences stabilizing coexistence and average fitness differences promoting competitive exclusion, the condition for coexistence (mutual invasibility) is expressed as (11, 36):

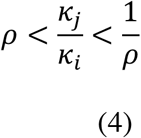

This condition shows that species with large differences in fitness need to also have high niche differences to coexist. In contrast, species with similar fitness can coexist even with small niche differences. As a consequence, the mutual invasibility criterion allows us to quantify the degree to which a pair of species can stably coexist. Species pairs whose niche differences are much larger than the minimum required to overcome the fitness differences between them will be more stable than species pairs whose niche differences are close to the minimum. Species pairs whose niche differences are smaller than the minimum needed to overcome fitness differences will be unstable. We used this condition to relate the degree of stability to productivity (see below *analyses* section).

### Field parameterization of population models under two contrasting climatic conditions

We conducted a field experiment to parameterize these models with estimates of species germination fractions, seed survival in the soil and per germinant fecundities in the absence of neighbors. We also estimated all pairwise interaction coefficients between the species by growing each species in competition with itself and with all other species, in experimental plant communities in which we manipulated competitor density and identity, following previous experimental designs (20). Specifically, we established 180 rectangular plots (0.65 m × 0.5 m) in September 2014 prior to the major autumn rains. We randomly assigned each of 80 plots to be sown with one of the 10 species at a density of 2, 4, 8, or 16 g/m^2^ of viable seed, giving two replicates per density and per species. Each plot was divided into 20 subplots (a four row by five column array) with a buffer of 2 cm along the edge of the plot. At the center of each subplot, we sowed five viable seeds of one of the 10 species, and germinants were thinned to a single individual per subplot. This design allowed us to measure viable seed production on two focal individuals per species and plot, when competing with different number of neighbors of the same species and each of the other 9 species. We additionally established 10 plots that had the same array but did not include any density treatment in order to measure viable seed production of focal individuals of the 10 species in the absence of competition. Information from both plots with and without density treatments were combined to estimate per germinant seed production in the absence of neighbors (*λ*_i_) and the interaction coefficients (*α*_ij_) according to the function (20).

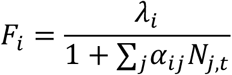

To fit this function, we used maximum likelihood approaches (optim method=“L-BFGS-B”) to ensure that *λ*_*i*_ and *α*_*ij*_ ≥ 0. For each target species *i*, we fit a separate model jointly evaluating its response to individuals of all other species and itself. This approach fits a single per germinant fecundity in the absence of competition, *λ*_*i*_ for each species *i*. To estimate species’ germination fraction (*gi*), we counted the number of germinants in at least one plot of each density and divided by the total number of seeds originally sown. To obtain the seed bank survival (*s*_*i*_), we followed the method detailed in (36), burying 5 replicates of 100 seeds each on the surrounding area from September 2014 to August 2015 and determining their viability as described in (7). Finally, we repeated the same experiment with the remaining 90 plots, sowing seeds on 10^th^ December 2014 to simulate a drier climate. We selected this type of treatment because annual species germination occurs only after major autumn rains and, in Mediterranean ecosystems, delays in the start of the rainy season strongly affect annual plant population dynamics (40). This delay of 64 days resulted in changes in daylight, temperature, and rainfall between treatments. However, most notably, it produced a 38.7% reduction in precipitation (from 532 in the first experiment to 326 mm for this second experiment).

### A biodiversity-functioning experiment with multiple functions

We conducted a biodiversity-functioning experiment to simultaneously estimate complementarity and selection effects for three different functions: biomass production, litter decomposition, and changes in soil nutrient content. We established 104 circular plots (0.75m^2^) in the same area and at the same times as the competition experiment. We randomly assigned each plot to be a monoculture or a mixture of 3, 5, 7, 9 and 10 species. All plots were sown at a total seed density of 15g/m^2^, and seed mass was evenly divided between the species in mixtures. To create the mixtures, we randomly assembled 6 different communities of 3 species, 4 communities of 5 species, 3 communities of 7 species, and 2 communities of 9 species. These communities, as well as the 10 monocultures and the one 10 species mixture, were all replicated twice within each climatic condition (i.e. climate control and drought). We visually assessed the biomass of each plot biweekly, and collected aboveground biomass when it was maximal in each plot. We defined the peak of biomass as the first date when a majority of species were senescent. At this time, all species had produced flowers. Biomass was separated by species, air dried for two weeks, then oven dried at 60 °C during 3 days and weighed (g).

In addition, we conducted biweekly surveys of leaf senescence within species to estimate when to put litter bags in the soil. We defined the peak of leaf senescence as the date when the number of individuals with clear senescence symptoms (several leaves dropped from the individuals) outnumbered those without. Litter bags initially contained between 0.35 and 1.5 g of leaf litter material from a single species, which was collected from individuals of the same plot where we placed the bags. This procedure ensures that litter quality and litter decomposition rates are driven by the species characteristic and by the specific competitive, soil, and microenvironmental conditions of each plot. We separated litter bags for each of the species included in the plot. This might underestimate litter mixing effects but the alternative, a single litter bag with mixed litter, would not have allowed us to distinguish the identity of decomposed litter and therefore to estimate decomposition rates at the species level. After three months, litter bags were harvested, carefully brushed clean, dried at 60 °C during 3 days, and weighed to calculate the percentage of litter mass loss.

We assessed soil nutrient dynamics as changes in C, N, P, and K, Ca, Mg cations right before (September 2014) and after the experiment (September 2015), in the first 10 cm of soil. This corresponds to the soil depth influenced by annual plant vegetation in Mediterranean ecosystems and contains 95% of the total community root biomass (41). For chemical analyses, soils were dried in the lab at 30°C until constant weight, and sieved (2mm) to eliminate stones and large roots. Soils were analyzed for total organic C (%) (Walkley-Black method (42)), total organic N (%) (Kjeldahl method (43)), available P (mg/kg) (Olsen method (44), and exchange cations (mg/kg) (Ca^2+^, Mg^2+^, K^+^, extracted with 1M ammonium acetate and determined by atomic absorption).

### Analyses

We first explored the relationships between species diversity and biomass production, litter decomposition and nutrient content at the end of the experiment. We tested for linear and non-linear saturating relationships for the three types of functions.

Then, we tested for correlations between complementarity/selection effects and niche/fitness differences, under the two climatic conditions and for the three functions considered. Because niche and fitness differences are defined as pairwise measures, we could not use the standard additive partitioning approach to calculate them (10) and instead we used diversity interaction models (25) to calculate complementarity between pairs of species and selection effects for individual species. These models estimate selection effects as the ability of species to dominate each function regardless of its initial abundance because possess particular traits such as high leaf N or high photosynthetic rates. On the other hand, complementarity occurs when species yield more function in mixture than in monoculture because for instance they partition resource use. In order to convert selection effects to a pairwise measure we calculated the ratio between selection effects for pairs of species. We used the ratio rather than a difference because fitness differences are also defined as a ratio between species fitnesses (see equation 3). We then checked whether the selection and complementarity effects from the diversity interaction models (25) correlated with those produced by the additive partition of Loreau and Hector (10). In order to do this, we summed the individual (selection), or pairwise (complementarity) values from the diversity interaction models across all species in each community. These values correlated reasonably well with the values from the additive partitioning (r-values ranging between 0.487-0.769; *SI Appendix*, Fig. S2).

We used Mantel tests, and the Benjamini and Hochberg correction for multiple comparisons, to test for significant correlations between coexistence (niche and fitness differences, equations 2 and 3) and biodiversity functioning mechanisms (complementarity and selection effects). In addition to analysing the overall fitness differences we also split them into their two components, the demographic ratio and the response to competition ratio, and correlated each component with complementarity and selection effects. The same Mantel test procedure was also used to test for the correlation between the stability of species pairs (difference between the observed niche difference and the minimum niche difference needed to allow coexistence) and the degree of function predicted for that pair. We used our diversity interaction models to estimate the degree of functioning predicted for each species pair.

Finally, we combined field-parameterized models of competition among 10 annual plant species with tools from network theory to analyze whether there are feasible and locally stable equilibrium for any of the 32 experimentally assembled communities. More specifically, we first used an algebra matrix inversion approach (45) to compute species abundances at a single stable equilibrium for these combinations of 3, 5, 7, 9, and 10 species. Because species abundance at equilibrium can be positive or negative, this means that a stable equilibrium might not contain all species from the original community pool. Therefore, we estimated which of the stable species assemblages, were also feasible, i.e. assemblages with positive abundances of all members at equilibrium, by deriving the Jacobian matrix for the annual plant model, and assessing whether the maximal eigenvalue of the Jacobian was less than one, in absolute terms (46). This network analysis was used to predict whether the observed levels of the different ecosystem functions were likely to be stable over time, or transient, based on which species were expected to go extinct. All analyses were conducted in R Version. 3.5.3 (47).

## Acknowledgements

We thank Eduardo Gutierrez and Juan S. Cara for conducting soil nutrient analyses and lab guidance. O. G. acknowledges postdoctoral financial support provided by the European Union Horizon 2020 research and innovation program under the Marie Sklodowska-Curie grant agreement No 661118-BioFUNC.

## Supplementary material

**Fig. S1.**
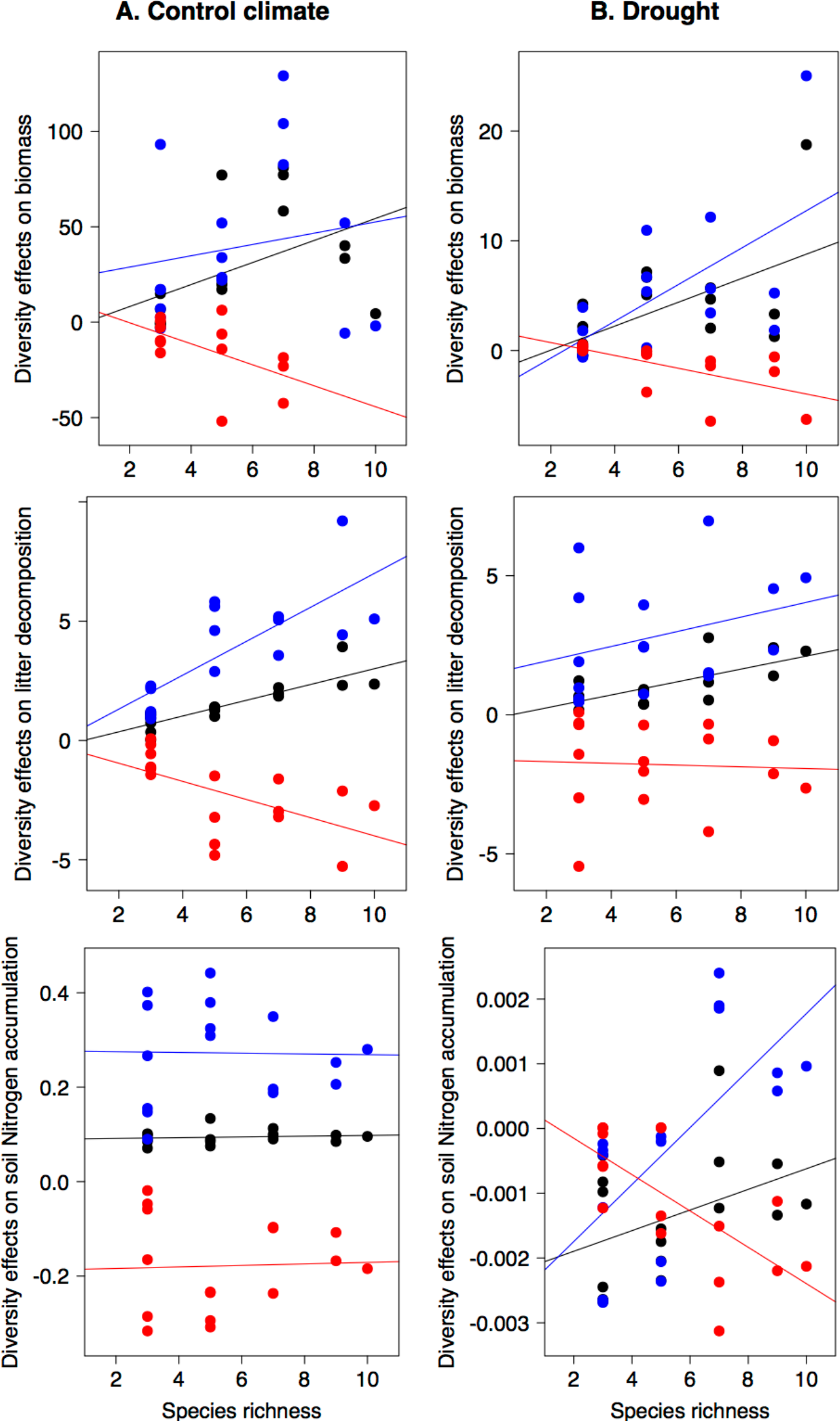
Net biodiversity effects (black line) and its two components (complementarity (blue line) and selection effects (red line)) as a function of species richness across the three functions considered (biomass production, litter decomposition, and soil nutrient accumulation under the two contrasted climatic conditions (A) Control climate and (B) Drought. Species richness was a significant predictor (p<0.05) of the net effect of biodiversity on productivity for all functions except for litter decomposition under drought conditions and soil nitrogen under control climate. For soil nutrients, we represent the particular case of nitrogen but very similar relationships were observed for the other elements considered.

**Fig. S2.**
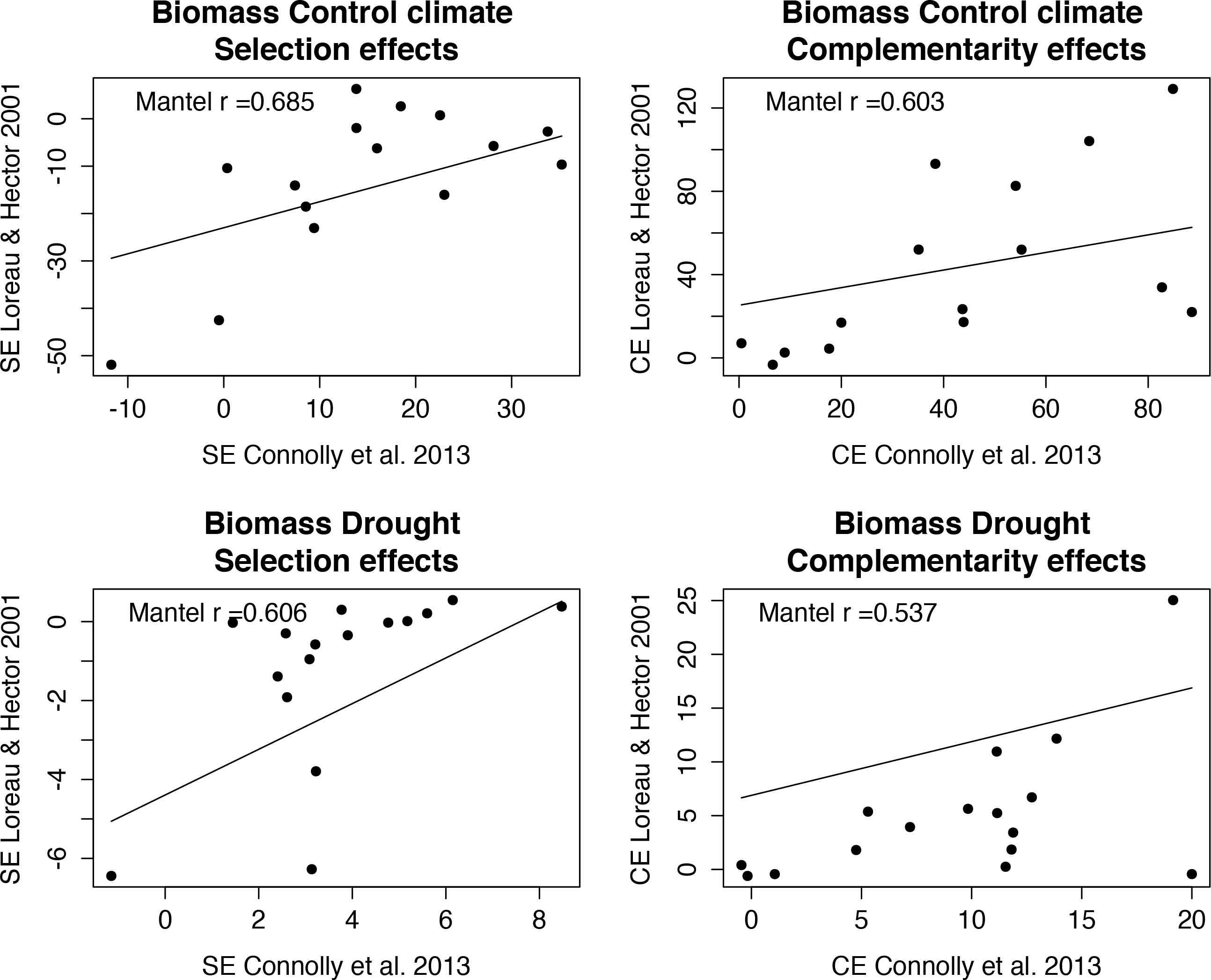
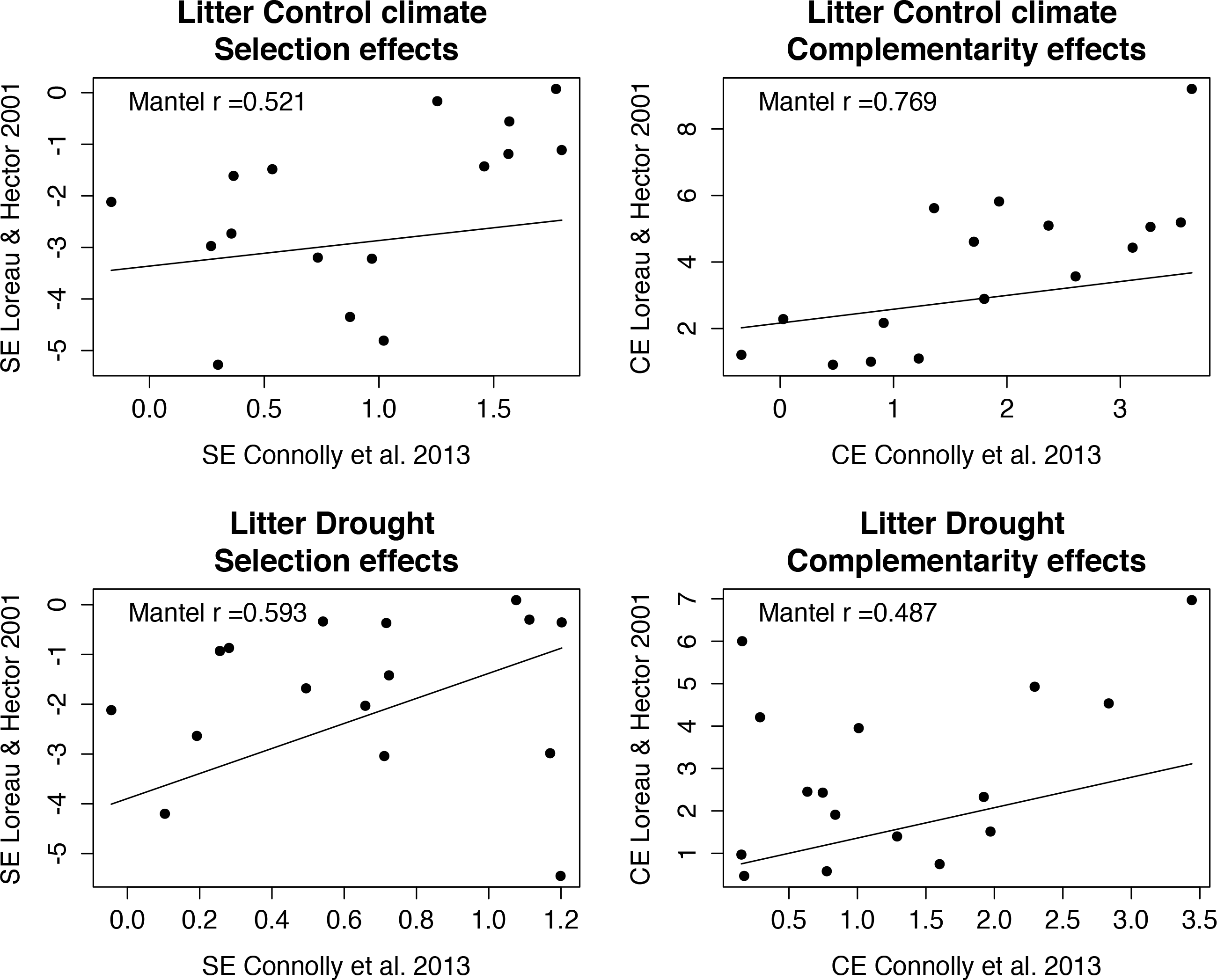
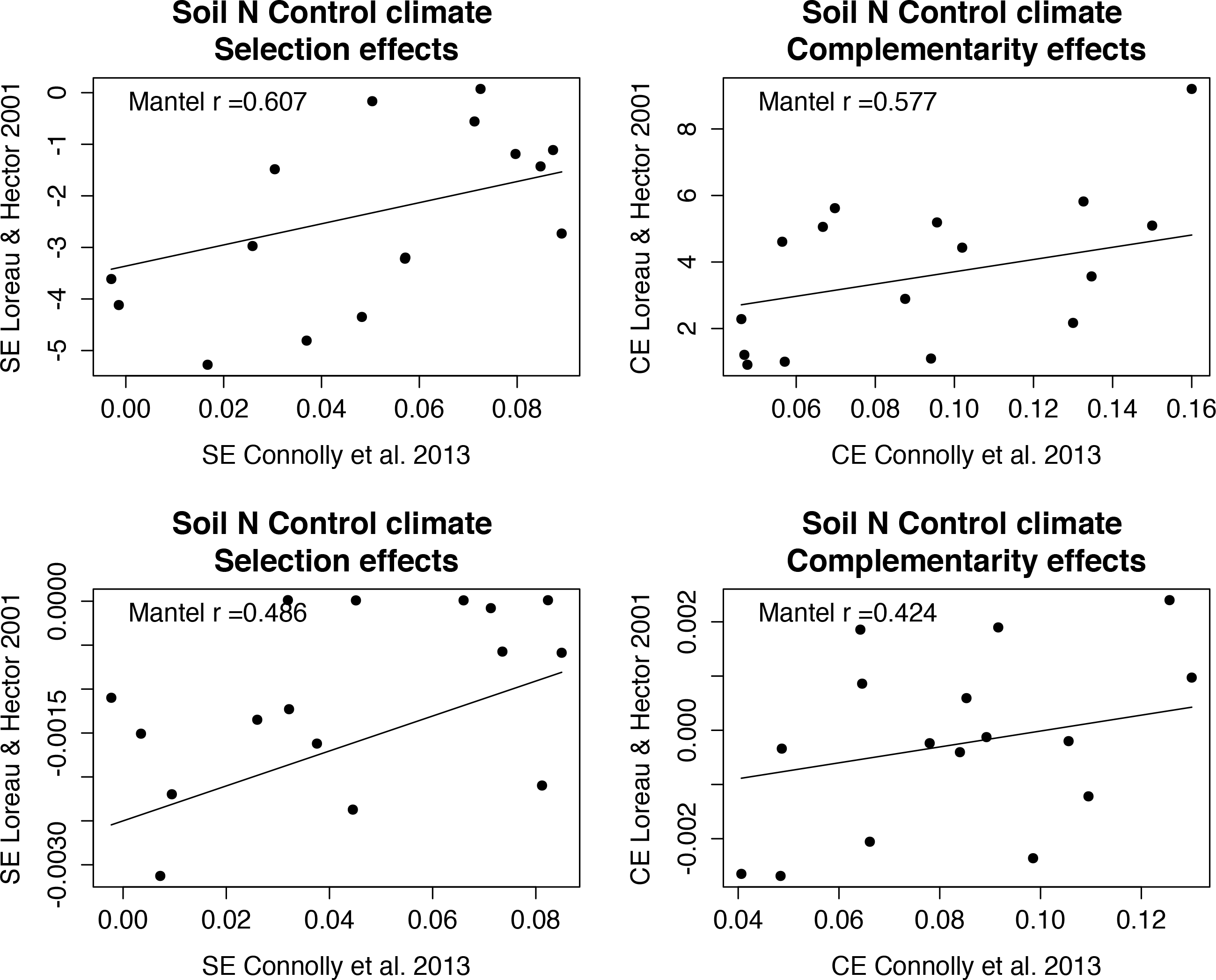
Correlation between complementarity effects and differences in species’ selection effects obtained from additive partitioning (10) (y-axis) and from diversity interaction models (25) (x-axis). These graphs show that for the multiple functions (biomass, litter decomposition, soil nutrient content) evaluated in our experiment complementarity and selection effects correlated well between the two approaches under control climate (upper panels) and drought conditions (bottom panels). To obtain complementarity and selection effects at the community level using diversity interaction models (25), we sum all complementarity effects between the species in the community and all selection effects for the individual species.

**Fig. S3.**
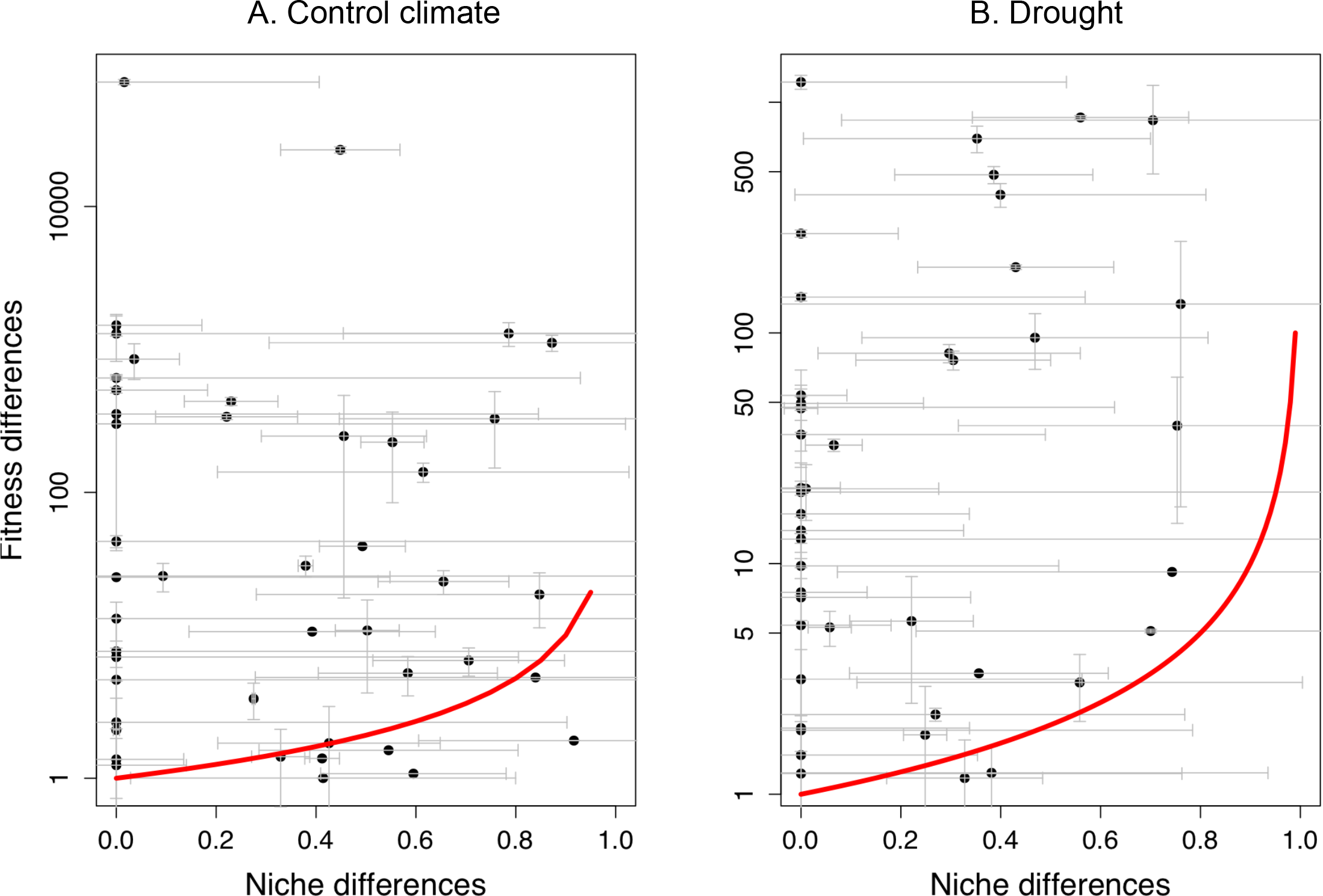
Relationship between pairwise niche and fitness differences for the two climate scenarios: (A) control climatic scenario for the study area where the first major rains after the drought period occurred in October, and (B) drought event where first major rains came in December, causing a 2-month delay in seed germination. Note that each point is a pair of species, and error bars are shown for niche and fitness differences. Also note, the differences in magnitude of the y-axis between climatic treatments. In fact, drought significantly reduced fitness differences and marginally reduced niche differences (paired t-test, *t* = 2.3564, *df* = 44, *P* = 0.0230 and *t* = 1.7493, *df =* 44, *P* = 0.0872, respectively)., The red solid line separates the region where the condition for coexistence is met (ρ< κ_j_/κ_i_) from the competitive exclusion region. Six species pairs fall in the coexistence region under control climatic conditions and two under drought. For the rest of the species pairs, average fitness differences exceed stabilising niche differences. Note that our experiment focused on interactions at the neighbourhood spatial scale over a single generation and therefore does not capture the spatial and temporal heterogeneity that allows these pairs to coexist at the landscape scale.

**Fig. S4.**
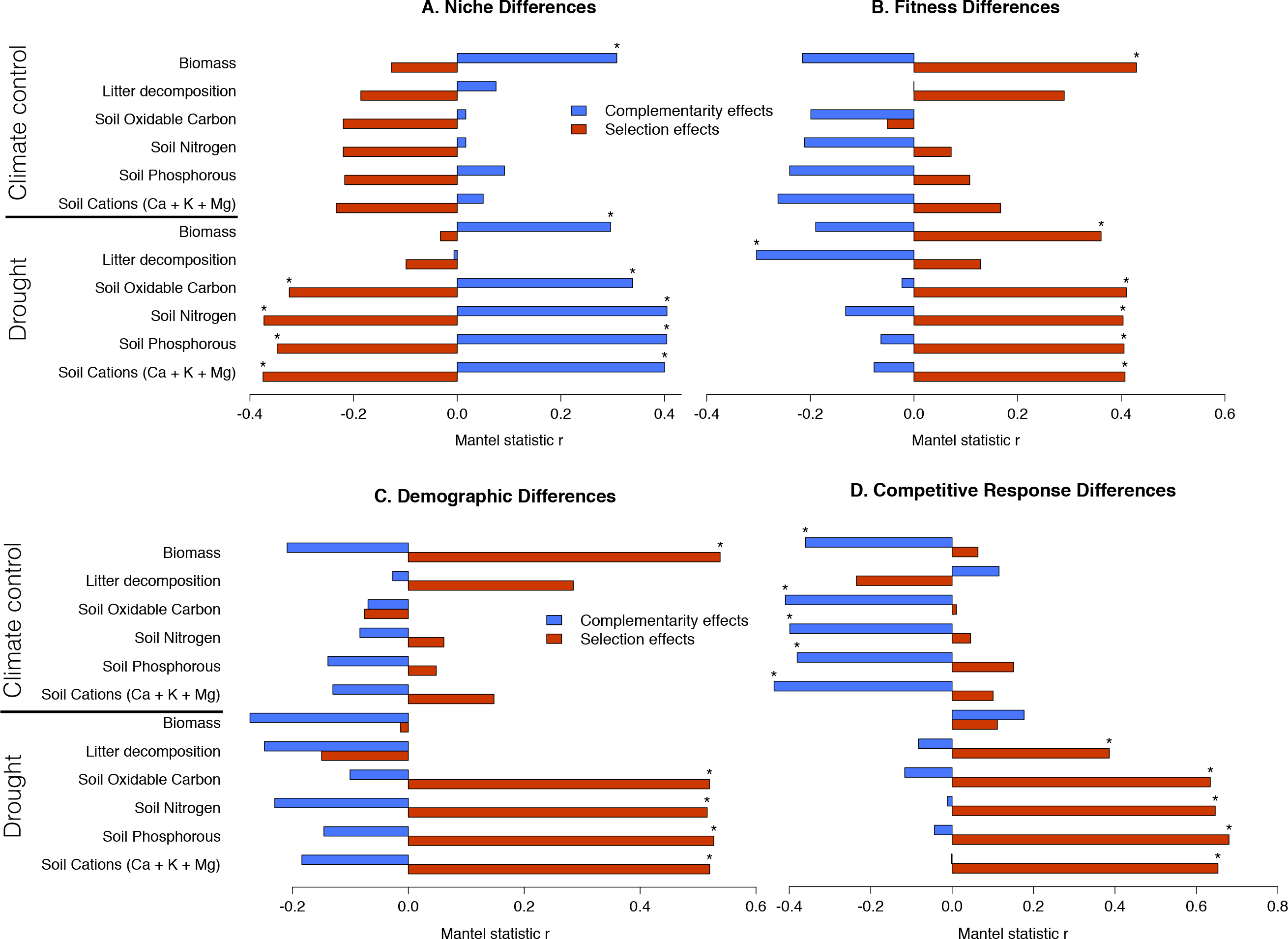
Same as Figure 2 in the main text, this supplementary figure shows correlations between (A) stabilizing niche and (B) average fitness differences and complementarity (blue) and selection effects (red), here for all soil nutrients including soil oxidable C, soil P, and soil cations. Correlations between complementarity and selection are also shown with the two components of fitness differences, the demographic ratio (C) and the competitive response ratio (D). Soil oxidable C, soil P, and soil cations. Correlations are shown for the two climates (control and drought). Significant correlations, following Benjamini-Hochberg correction for multiple comparisons, are marked with an asterisk.

**Fig. S5.**
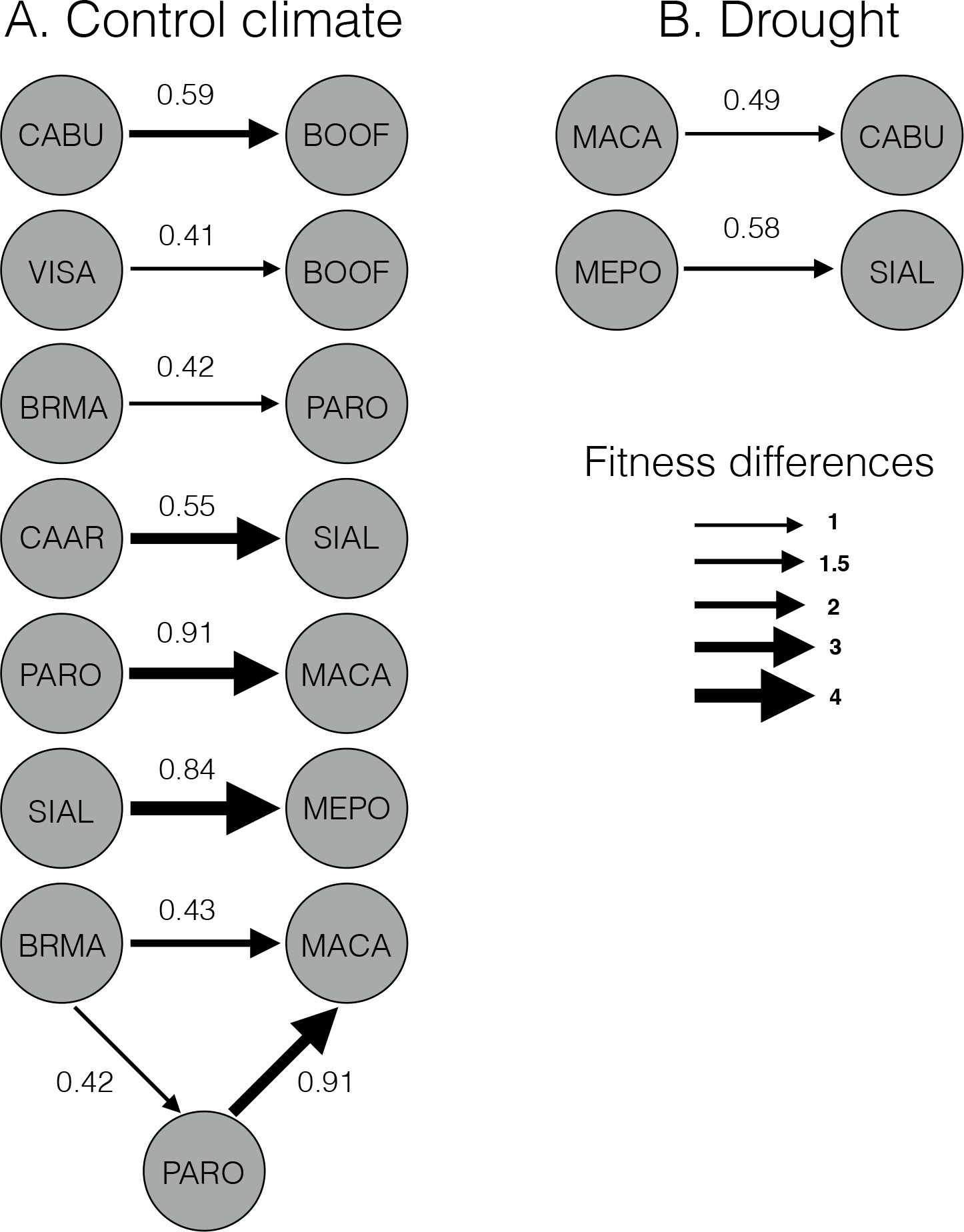
Illustration of the 6 species pairs and one triplet that produced a feasible and stable equilibrium under control climatic conditions (A), and the 2 species pairs with the same properties under drought (B). Note that none of the experimentally assembled communities in which we estimated the effect of species diversity on ecosystem functioning (see Table 1 for communities and species code) are predicted to stably coexist. Therefore, the biodiversity effects found in our experiment were transient rather than stable, which means that the degree of function is predicted to change with time as species are competitively excluded from communities, see Fig. S5. Black arrows denote the magnitude of fitness differences, and their direction indicates the best competitor within species pairs or triplets. Pairwise niche differences (between 0 and 1) are provided numerically for each species pair.

## References

1. S. Soliveres, et al., Biodiversity at multiple trophic levels is needed for ecosystem multifunctionality. Nature 536, 456–459 (2016).

2. F.T. Maestre, et al., Plant species richness and ecosystem multifunctionality in global drylands. Science 335, 214–218 (2012).

3. E.S. Zavaleta, J. R. Pasari, K. B. Hulvey, D. Tilman, Sustaining multiple ecosystem functions in grassland communities requires higher biodiversity. Proc. Natl. Acad. Sci. U.S.A 107, 1443–1446 (2010).

4. O. Hoegh-Guldberg, et al., Coral reefs under rapid climate change and ocean acidification. Science 318, 1737–1742 (2007).

5. B. Worm, et al., Impacts of biodiversity loss on ocean ecosystem services. Science 314, 787–790 (2006).

6. P. B. Adler, H. J. Dalgleish, S. P. Ellner, Forecasting plant community impacts of climate variability and change: when do competitive interactions matter? J. Ecol., 100, 478–487 (2012).

7. L. Matias, O. Godoy, L. Gómez-Aparicio, I. M. Pérez-Ramos, An experimental extreme drought reduces the likelihood of species to coexist despite increasing intransitivity in competitive networks. J. Ecol. 106, 826–837 (2018).

8. T. Newbold, et al., Global effects of land use on local terrestrial biodiversity. Nature 520, 45 (2015).

9. E. Allan, et al., Land use intensification alters ecosystem multifunctionality via loss of biodiversity and changes to functional composition. Ecol. Lett., 18, 834–843 (2015).

10. M. Loreau, A. Hector, Partitioning selection and complementarity in biodiversity experiments. Nature 412, 72–76 (2001).

11. P. Chesson, Mechanisms of maintenance of species diversity. Annu. Rev. Ecol. Syst. 31, 343–366 (2000).

12. K. E. Barry et al., The future of complementarity: disentangling causes from consequences. Trends Ecol. Evol. 34, 167–180 (2018).

13. J. W. Fox, The long-term relationship between plant diversity and total plant biomass depends on the mechanism maintaining diversity. Oikos 102, 630–640 (2003).

14. D. Tilman, C. L. Lehman, K. T. Thomson, Plant diversity and ecosystem productivity: theoretical considerations. Proc. Natl. Acad. Sci. U.S.A 94, 1857–1861 (1997).

15. L. A. Turnbull, J. M. Levine, M. Loreau M, A. Hector, Coexistence, niches and biodiversity effects on ecosystem functioning. Ecol. Lett. 16, 116–127 (2013).

16. L. A. Turnbull, F. Isbell, D.W. Purves, M. Loreau M, A. Hector, Understanding the value of plant diversity for ecosystem functioning through niche theory. Proc. R. Soc. B p 20160536. (2016)

17. P. J. Vermeulen, J. Ruijven, N. P. R. Anten, W. Werf, A. Satake, An evolutionary game theoretical model shows the limitations of the additive partitioning method for interpreting biodiversity experiments. J. Ecol. 105, 345–353 (2017).

18. I. T. Carroll, B. J. Cardinale, R. M. Nisbet, Niche and fitness differences relate the maintenance of diversity to ecosystem function. Ecology 92,1157–1165 (2011).

19. H. Maherali, J. N. Klironomos, Influence of phylogeny on fungal community assembly and ecosystem functioning. Science 316, 1746–1748 (2007).

20. O. Godoy, N. J. B. Kraft, J. M. Levine, Phylogenetic relatedness and the determinants of competitive outcomes. Ecol. Lett. 17, 836–844 (2014).

21. J. M. Levine, J. HilleRisLambers, The importance of niches for the maintenance of species diversity. Nature 461, 254–257(2009).

22. D. Craven, et al., Plant diversity effects on grassland productivity are robust to both nutrient enrichment and drought. Phil. Trans. R. Soc. B 371, 20150277 (2016).

23. N. R. Guerrero-Ramírez, et al., Diversity-dependent temporal divergence of ecosystem functioning in experimental ecosystems. Nature Ecol. Evol. 1, 1639 (2017).

24. F. Isbell, et al., Quantifying effects of biodiversity on ecosystem functioning across times and places. Ecol. Lett. 21, 763–778 (2018).

25. J. Connolly et al., An improved model to predict the effects of changing biodiversity levels on ecosystem function. J. Ecol. 101, 344–355 (2013).

26. I. J. Wright et al., The worldwide leaf economics spectrum. Nature 428, 6985 (2004).

27. N. DeMalach, E. Zaady, R. Kadmon, Light asymmetry explans the effect of nutrient enrichment on grassland diversity. Ecol. Lett. 20, 60–69 (2017).

28. S. Hättenschwiler, A.V. Tiunov, S. Scheu, Biodiversity and litter decomposition in terrestrial ecosystems. Annu. Rev. Ecol. Evol. Syst. 36, 191–218 (2005).

29. O. Godoy, I. Bartomeus, R. Rohr, S. Saavedra, Towards the integration of niche and network theories Trends Ecol. Evol 33287–300 (2018).

30. U. Brose, H. Hillebrand, Biodiversity and ecosystem functioning in dynamic landscapes. Proc. R. Soc. B 371(2016).

31. I. M. Pérez-Ramos, et al., Climate variability and community stability in Mediterranean shrublands: the role of functional diversity and soil environment. J. Ecol. 105, 1335–1346 (2017).

32. B J. Cardinale, M. A. Palmer, S. L. Collins, Species diversity enhances ecosystem functioning through interspecific facilitation. Nature 415, 426–429 (2002).

33. L. Wittebolle, et al., Initial community evenness favours functionality under selective stress. Nature 458, 623–626 (2009).

34. J. M. Levine, J. Bascompte, P. B. Adler, S. Allesina, Beyond pairwise mechanisms of species coexistence in complex communities. Nature 546, 56–64 (2017).

35. S. Saavedra, et al., A structural approach for understanding multispecies coexistence. Ecol. Monogr. 87, 470–486 (2017).

36. O. Godoy, J. M. Levine, Phenology effects on invasion success: insights from coupling field experiments to coexistence theory. Ecology 95, 726–736 (2014).

37. P. L. Chesson, Geometry, Heterogeneity and Competition in Variable Environments. Proc. R. Soc. B 330, 165–173 (1990)

38. I. M. Pérez-Ramos, et al., Functional traits and phenotypic plasticity modulates species coexistence across climatic conditions. Nat. Comm. 10, 2555 (2018).

39. J. B. Lanuza, I. Bartomeus, O. Godoy, Opposing effects of floral visitors and soil conditions on the determinants of competitive outcomes maintain species diversity in heterogeneous landscapes. Ecol. Lett. 21, 865–874 (2018).

40. J. M. Levine, A. K. McEachern AK, C. Cowan, Seasonal timing of first rain storms affects rare plant population dynamics. Ecology 92, 2236–2247 (2011).

41. M. Rueda, S. Rebollo, M. A. Rodríguez, Habitat productivity influences root mass vertical distribution in grazed Mediterranean ecosystems. Acta Oecologica 36, 377–382 (2010).

42. A. Walkley, I. A. Black, An examination of the Degtjareff method for determining soil organic matter, and a proposed modification of the chromic acid titration method. Soil Science 37, 29–38. (1934)

43. J. Bremner, D. R. Keeney, Steam distillation methods for determination of ammonium, nitrate and nitrite. Analytica chimica acta 32, 485–495 (1965).

44. S. R. Olsen, Estimation of available phosphorus in soils by extraction with sodium bicarbonate (USDA; Washington) (1954).

45. P. Yodzis, The indeterminacy of ecological interactions as perceived through perturbation experiments. Ecology 69, 508–515 (1988).

46. O. Godoy, D. B. Stouffer, N. J. B. Kraft, J. M. Levine, Intransitivity is infrequent and fails to promote annual plant coexistence without pairwise niche differences. Ecology 98,1193–1200 (2017).

47. R Development Core Team R: A language and environment for statistical computing (R Foundation for Statistical Computing, Vienna, Austria), 3.5.3. (2019).

